# From Structure to Dynamics: Activation Mechanism of the G Protein-Coupled Bile Acid Receptor 1-G*s* Complex

**DOI:** 10.64898/2026.02.06.704396

**Authors:** Bianca Fiorillo, Federica Moraca, Francesco Saverio Di Leva, Valentina Sepe, Stefano Fiorucci, Vittorio Limongelli, Angela Zampella, Bruno Catalanotti

**Author notes:** To whom correspondence should be addressed. Bianca Fiorillo Bruno Catalanotti.

## Abstract

The G protein-coupled bile acid receptor 1 (GPBAR1, also known as TGR5) is a key mediator of bile acid signaling, exerting its physiological effects through coupling with the stimulatory G protein (G*_s_*). This interaction is essential for stabilizing the receptor’s active conformation and triggering downstream signaling.

Among endogenous ligands, lithocholic acid (LCA) is the most potent natural agonist. However, the dynamic features underlying its binding and activation mechanisms remain poorly defined.

In this study, we investigated the molecular basis of the interaction between LCA and GPBAR1, as well as the functional consequences of this interaction on receptor activation by integrating homology modelling, molecular docking, and molecular dynamics (MD) simulations.

Our calculations reveal that LCA binding stabilizes the active state of GPBAR1, biasing the conformational ensemble of TM5 and TM6, as well as the main microswitches. These ligand-induced rearrangements enhance the coupling interface with the α5 helix of Gα*_s_* and facilitate allosteric communication between the orthosteric and intracellular sites.

Overall, our findings provide dynamic insight into how LCA modulates GPBAR1 activation and G protein engagement, highlighting its role as a molecular effector in bile acid signaling, and furnishing molecular detail relevant to ongoing efforts in GPBAR1-targeted compound development.

## 1. Introduction

The G protein-coupled bile acid receptor 1 (GPBAR1), also known as TGR5, is a class A G protein-coupled receptor (GPCR) that acts as a membrane-bound sensor for bile acids, integrating metabolic and immune signaling across multiple physiological systems. GPBAR1 is expressed in a variety of tissues, including enteroendocrine cells, hepatocytes, brown adipose tissue, skeletal muscle, and immune cells such as macrophages and monocytes. Upon activation by endogenous bile acids, GPBAR1 preferentially couples to the stimulatory G protein G*_s_*, leading to increased intracellular levels of cyclic AMP (cAMP) and subsequent activation of protein kinase A (PKA)-dependent pathways that regulate gene transcription and cellular function.[1,2] This receptor has emerged as a promising pharmacological target in recent years, due to its central role in modulating glucose and lipid metabolism, energy expenditure, intestinal hormone secretion, and inflammatory responses.[2–6]

In the gastrointestinal tract, the activation of GPBAR1 on enteroendocrine L cells promotes the release of incretin hormones, particularly glucagon-like peptide-1 (GLP-1), thereby enhancing insulin secretion and contributing to glycaemic control.[2,5,7,8] In brown adipose tissue and skeletal muscle, GPBAR1 signaling promotes mitochondrial activity and thermogenesis, supporting increased energy expenditure and lipid mobilization, which are beneficial in the context of obesity and type 2 diabetes. [9] Finally, in immune cells, particularly monocytes and macrophages, GPBAR1 activation has been shown to suppress pro-inflammatory cytokine production and shift macrophage polarization towards an anti-inflammatory (M2-like) phenotype, revealing potential for therapeutic modulation in chronic inflammatory conditions such as inflammatory bowel disease (IBD).[10,11]

Altered expression or signaling through GPBAR1 has been implicated in the pathogenesis of several diseases. Reduced receptor activity, for example, has been associated with insulin resistance and impaired glucose tolerance, while aberrant or constitutive activation may be involved in the progression of certain cancers, such as cholangiocarcinoma, where bile acid signaling intersects with oncogenic pathways.[6,12,13] These observations highlight the importance of gaining a detailed molecular understanding of GPBAR1 activation, ligand binding, and downstream effector coupling in both physiological and pathological contexts.

Endogenous GPBAR1 ligands represent a diverse family of bile acids, and it is their structural variability that underpins differential agonistic efficacy and tissue-selective pharmacology. Beyond their physiological roles, these compounds are increasingly recognized as privileged scaffolds for drug design, particularly in the treatment of metabolic and inflammatory disorders. Among endogenous bile acids, lithocholic acid (LCA) (Figure 1) is the most potent GPBAR1 agonist (EC_50_ = 0.53 μM),[14] displaying a high binding affinity and robust G*_s_*-mediated signaling. This makes LCA a benchmark ligand in both experimental and computational studies of GPBAR1.[15,16]

**Figure 1.**
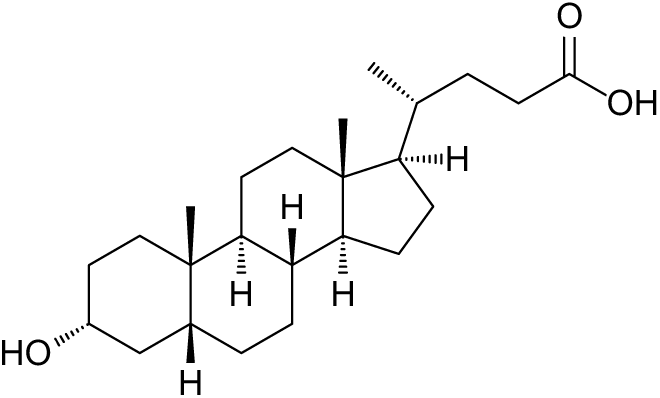
Two-dimensional (2D) chemical structure of LCA.

Until recently, the lack of experimentally determined GPBAR1 structures severely limited a mechanistic understanding of bile acid recognition and receptor activation. In this context, homology modelling and other computational approaches, together with structure–activity relationship (SAR) and site-directed mutagenesis studies, provided the primary framework for interpreting ligand binding and receptor activation.[17–19] Early models, such as that developed by D’Amore et al.,[20] exploited the conservation of class A GPCRs to propose active-like receptor conformations and rationalize bile acid binding modes, guiding docking, virtual screening, and mutagenesis experiments. In particular, these studies suggested that subtle differences in hydroxylation patterns and side-chain flexibility translate into distinct receptor engagement modes and downstream signaling biases.[21,22] and identified key residues within TM3, TM5, TM6, and TM7 as important for ligand recognition. However, in the absence of direct structural information, how different agonists stabilize specific active-state conformations and how these conformations couple to G*_s_* remained largely unresolved. A major breakthrough came with the publication of cryo-electron microscopy (cryo-EM) structures of GPBAR1 in complex with the G*_s_* protein, including the INT-777 bound assembly (PDB ID: 7CFN),[21] which offered the first high-resolution view of the active receptor-G protein complex. Nevertheless, these structures lack unresolved flexible loops, the terminal regions of GPBAR1 and G*α_s_*, and the entire α-helical domain (AHD) of Gα_s_, which is critical for nucleotide exchange. Furthermore, static cryo-EM models fail to capture the dynamic interplay between the receptor, the G protein, and the surrounding lipid bilayer. This issue has become even more pertinent in light of recent advances in GPCR medicinal chemistry, including the development of emerging molecular glues and allosteric modulators that act at the receptor-G protein interface to modulate transducer signaling,[23,24] which necessitate comprehensive, fully resolved structural representations of the entire activated macromolecular complex. To address these limitations, computational modelling is required to extend experimental structures into functionally complete and dynamically tractable systems.

In this study, we constructed and refined a full-length GPBAR1-Gα*_s_*β_1_γ_2_ model based on the INT-777-bound cryo-EM structure assembly (PDB ID: 7CFN),[21] embedded in a lipid bilayer and incorporating the native post-translational modification of N-linked glycosylation at Asn76^ECL1^ (superscripts refer to Ballesteros-Weinstein numbering),[25,26] to mimic a physiologically relevant membrane-associated state. Using this model, we investigated how the endogenous agonist LCA interacts with the orthosteric site and modulates the conformational landscape of GPBAR1 and its coupling to G_s_. By comparing LCA-bound and ligand-free microsecond-scale molecular dynamics simulations, we characterized how agonist binding reshapes receptor flexibility, microswitches organization, and long-range allosteric communication between the ligand-binding pocket and the intracellular G-protein interface.

This integrative computational framework provides a dynamic perspective on how endogenous ligands, such as LCA, modulate GPBAR1 signaling at the molecular level, prompting the rational design of synthetic agonists, allosteric modulators, or biased ligands with enhanced efficacy and selectivity for the treatment of metabolic and inflammatory diseases.

## 2. Material and Methods

### 2.1 Comparative modeling, structure refinement and system preparation

The cryo-EM structure of INT-777 bound GPBAR1-Gα*_s_*β_1_γ_2_ complex (PDB ID: 7CFN) [21] was selected as template for homology modeling. Prior to model building, nanobody fragments, the ligand, cholesterol, palmitic acid, and water molecules were removed. Missing segments in both the receptor and G protein subunits, which were unresolved in the experimental structure, were modelled using MODELLER version 10.6,[27] based on predicted coordinates retrieved from the AlphaFold database (https://alphafold.ebi.ac.uk).[28] In particular, the GPBAR1 sequence was aligned to the AlphaFold-predicted active-state structure, and the missing regions Ile13^N-ter^-Ala17^1.35^ and Thr288^8.51^-Ala306^C-ter^ were reconstructed. For the G_α_ subunit, the Met1-Lys8, Arg61-Thr204, and Asn254-Thr263 segments were derived from the AlphaFold-predicted model, after alignment on the Ras domain backbone. For G_β_, the N-terminal residue Met1 was modelled *ab initio*. Similarly, for G_γ_ the N-terminal (Met1-Asn4) and C-terminal (Glu63-Leu71) fragments were reconstructed *ab initio*. Predicted secondary structure elements were incorporated to refine poorly conserved regions, and the resulting models were ranked according to their discrete optimized protein energy (DOPE) score. The conformation with the highest score was selected and structurally validated via Ramachandran plot and MolProbity analysis,[29] showing >95% of residues in favored regions and Z-scores (-1.43 ± 0.24) comparable to experimentally solved GPCR-G*_s_* complexes. Importantly, the addition of the α-helical domain (AHD) of Gα*_s_* restored the canonical nucleotide-exchange architecture. The final full-length model thus provided a structurally coherent framework suitable for docking and MD studies. The resulting model was then processed using the Protein Preparation Wizard implemented in Maestro software (Schrödinger Suite 2022-4),[30] which involved the assignment of proper bond orders, the addition of hydrogens, the optimization of protonation states at physiological pH, and the minimization of the structure to relieve steric clashes.

### 2.2 Ligand preparation

The three-dimensional structure of lithocholic acid (LCA) was generated using the graphical user interface (GUI) of Maestro ver. 13.4.[30] The protonation state at pH 7.4 in water was calculated using the Epik module.[31] Finally, the ligand was minimized using the OPLS2005 force field[32] through 2,500 iteration steps of the Polak-Ribiere Conjugate Gradient (PRCG)[33] algorithm.

### 2.3 Docking Calculations

Docking of LCA to the GPBAR1 receptor was performed using the Standard Precision (SP) docking protocol implemented in the Glide software package [34] and the OPLS 2005 force field.[32] A grid box of 3 × 3 × 3 nm was defined around the ligand-binding cavity of the GPBAR1 receptor. A total of 100 ligand conformations were generated within the orthosteric binding site of GPBAR1, and conformational sampling of the ligand was enhanced by a factor of 4. All generated poses were analyzed, and the one with the lowest energy values in both Glide Emodel and GlideScore was selected.[35] The selected pose exhibited a docking score of -6.691 kcal/mol and an orientation compatible with that observed for INT-777 in the experimental structure (RMSD = 0.08 nm, computed on the steroidal scaffold) (PDB ID: 7CFN).[21]

### 2.4 MD calculations

MD simulations were performed with GROMACS suite ver. 2021.5,[36] using the Amber ff14SB, Lipid 17, and the CHARMM General Force Field (CGenFF) parameters for the proteins, lipids, and ligand, respectively.[37–39] In both the ligand-free and the LCA-bound-GPBAR1-G protein systems, the GPCR was embedded into a heterogeneous bilayer, of sizes 16 x 16 nm, composed of 1-palmitoyl-2-oleoyl-sn-glycero-3-phosphocholine (POPC) and cholesterol (CHL) at a molar ratio of 7:3, using the Membrane Builder tool of the CHARMM-GUI web server (https://charmm-gui.org).[40,41] Prior to membrane generation, the proteins were reoriented by alignment with the corresponding OPM-deposited structure (https://opm.phar.umich.edu). [42] An N-linked glycosylation consisting of one N-acetylglucosamine (GlcNAc) moiety was added at Asn76^ECL1^.[26] Additionally, a disulfide bond was introduced between Cys85^3.25^ and Cys155^ECL2^(superscripts refer to Ballesteros-Weinstein numbering),[25] in accordance with known GPBAR1 structural features. Both the N- and C-termini of the protein chains were capped at the points of truncation to mimic the native terminal environments. The resulting system was solvated with explicit TIP3P water and 0.150 mM NaCl in a 13 × 13 × 17 nm box.

Energy minimization was performed using the steepest descent algorithm in a single-step procedure, with restraints gradually released.

The system was equilibrated using a multi-step protocol consisting of 16 sequential stages, designed to ensure a gradual relaxation of the structure prior to production simulations. Thermalization was carried out under an NVT ensemble by progressively heating the system from 0 K to 300 K using the Berendsen thermostat with a coupling constant of 0.1 ps. [43] Equilibration steps were performed under NPT conditions employing the Berendsen barostat with semi-isotropic pressure coupling at 1 bar, allowing the system density and simulation box dimensions to stabilize.

The leap-frog integration algorithm was used throughout, with all bonds involving hydrogen atoms constrained using the LINCS algorithm. [44,45] A timestep of 1 fs was employed during the initial heating phases and increased to 2 fs during the later equilibration stages. The total equilibration time amounted to approximately 18 ns, after which the system was considered stable in terms of temperature, pressure, and density.

Production simulations were then performed in the NPT ensemble using a timestep of 2 fs. Temperature was maintained at 300 K using the Nosé–Hoover thermostat, [46] while pressure was controlled at 1 bar with the Parrinello–Rahman barostat under semi-isotropic coupling. [47] Periodic boundary conditions (PBC) were applied in all directions, and long-range electrostatic interactions were treated using the particle-mesh Ewald method. Each production run was carried out for 500 ns.

Analysis of MD simulations was performed on the merged trajectory. Trajectory clustering was performed using the GROMACS gmx cluster tool with the GROMOS method,[48] employing a 0.3 nm cutoff and with a stride of 100. Root Mean Square Deviation (RMSD), Root Mean Square Fluctuation (RMSF) and the radius of gyration were calculated with GROMACS gmx rmsd, rmsf, and gyrate tools, respectively.[36,44] Finally, distances and microswitches dynamics were monitored with the MDanalysis tool. [49,50]

### 2.5 SIFt analyses

SIFt analyses [51] were carried out on the merged MD trajectories using a Python implementation. Ligand–receptor interactions involving both backbone and side-chain atoms were encoded as a 9-bit fingerprint capturing the following interaction types: hydrogen bonds with the protein acting as donor (Hbond_ProD) or acceptor (Hbond_ProA); apolar contacts defined by carbon–carbon proximity; aromatic interactions in face-to-face (Aro_F2F) and edge-to-face (Aro_E2F) geometries; electrostatic interactions with positively (Elec_ProP) or negatively charged residues (Elec_ProN). Apolar interactions were identified using a distance cutoff of 4.5 Å, aromatic and electrostatic interactions using a cutoff of 4.0 Å, and hydrogen bonds using a cutoff of 3.5 Å. The probability of each interaction was estimated within a two-state Markov framework by sampling the posterior distribution of the transition matrix with standard Dirichlet priors, as previously described in Noè et *al*. [52] The resulting interaction fingerprints were visualized and plotted using RStudio vers. 2025.05.1.

### 2.6 PCA Analysis

Principal component analysis (PCA)[53] of the GPBAR1-G protein systems in both the ligand-free and LCA-bound states was performed on TM5 and TM6 to investigate their collective motions. This analysis was carried out using gmx covar and gmx anaeig, after the trajectories were aligned to the receptor backbone and the analysis was restricted to the residues forming TM5 and TM6.[36,44] Prior to the calculations, the 1.5 μs MD trajectories of each system were stripped of solvent and ions, and global translational and rotational motions were then removed by fitting the trajectories to the receptor backbone of the first MD frame. This procedure allowed us to generate an average protein structure for each system, which was then employed as the reference structure for the PCA. The coordinate covariance matrix was generated and diagonalized to obtain the principal components (PCs) as eigenvectors and eigenvalues. The first four PCs were analyzed in detail. The dominant mode (eigenvector 1) describing the motions of TM5 and TM6 was subsequently visualized in PyMOL vers. 2.3.0, where the displacement vectors illustrate the magnitude and direction of the collective motions (see Movie 1).

### 2.7 Network-based analysis

Allosteric communication within GPBAR1 was analyzed using the MDpath software package,[54] which identifies the most probable routes of dynamic information transfer between residues by combining atomic correlation analyses with graph-theoretical approaches. For each simulation, the receptor was extracted from the trajectories, and a residue-residue network was constructed based on the normalized mutual information (NMI) of atomic fluctuations. To ensure a consistent comparison between the two states, the residues defined as “ligand-interacting” were selected from the centroid structure of the most populated receptor cluster obtained from the holo MD trajectory. These residues were in direct contact with LCA during the simulation’s most representative conformational state and they were then used as the starting points of the allosteric network in both the holo and ligand-free analyses.

The MDpath algorithm evaluates correlations between residues and constructs possible allosteric routes according to user-defined geometric and statistical parameters. In our analysis, a distance cutoff of 0.6 nm was applied to define residue connectivity within the graph (graphdist flag), while 0.3 nm and 1.2 nm were used as the minimum and maximum thresholds for contact formation (closedist and fardist, respectively).

To smooth the resulting paths, a spline factor of 0.2 was adopted (spline flag), and 200 bootstrap resamples were performed (bs flag) to estimate the statistical confidence of each pathway. The top 30 most probable paths (numpath) were retained for further analysis and visualization.

To accelerate the computation of correlation matrices and clustering steps, each trajectory was processed using 16 CPU cores (cpu flag). The resulting NMI matrices were clustered and the pathways with the highest confidence were extracted and ranked according to their occurrence probability.

The identified communication routes were finally mapped onto the receptor structure and visualized using PyMOL vers. 2.3.0 by connecting the Cα atoms of sequential residues along each path. This analysis identified the major allosteric communication networks connecting the ligand-binding pocket to the intracellular domain and revealed how ligand binding reshapes the internal signaling architecture of GPBAR1. Path weight values for each residue were calculated from MDPath results. For visualization, only residues whose cumulative path weight exceeded 0.5 were included, and the values were plotted as side-by-side bars for the ligand-bound and ligand-free systems using RStudio vers. 2025.05.1 and the ggplot2 package, with residue labels ordered according to the protein sequence.

MD trajectories were visualized using VMD software,[55] and all figures were rendered by UCSF Chimera [56] and PyMOL vers. 2.3.0 (https://pymol.org).

## 3. Results

### 3.1 A Structurally Validated Model of the Full-Length GPBAR1-G*_s_* Complex

A full-length model of the GPBAR1–Gα*_s_*β_1_γ_2_ complex was generated using MODELLER v10.6,[27] based on the cryo-EM structure of INT-777-bound GPBAR1-Gα*_s_*β_1_γ_2_ assembly (PDB ID: 7CFN). [21] This template was selected over the available AlphaFold models to ensure an active, ligand-bound receptor conformation related to our compound of interest.

The obtained model displays stereochemical quality comparable to that of experimentally determined GPCR–G*_s_* protein assemblies. Structural validation of the receptor indicated excellent geometry, with more than 95% of residues located in favored regions of the Ramachandran plot and MolProbity Z-scores (−1.43 ± 0.24) consistent with high-resolution cryo-EM structures.[29]

Importantly, the reconstructed model includes all previously unresolved regions of both the receptor and the G protein, particularly the α-helical domain (AHD) of Gα*_s_*, restoring the canonical architecture associated with nucleotide exchange and G*_s_* coupling, yielding a complete and coherent representation of the active signaling complex.

Overall, the resulting full-length GPBAR1–Gα*_s_*β_1_γ_2_ model provided a structurally consistent and functionally relevant framework for investigating ligand binding, receptor activation, and G protein engagement in subsequent computational studies.

### 3.2 The Dynamic Binding Mode of LCA Reveals Novel Stabilizing Interactions

To evaluate the binding mode of LCA at the GPBAR1 orthosteric site, we initially performed docking calculations with Glide.[30]

The top-ranked docking pose (-6.69 kcal/mol) places the steroidal core of LCA deep within the transmembrane bundle (Figure 2), establishing hydrophobic interactions with residues from TM2, TM3, TM6 and TM7, such as Leu71^2.60^, Leu74^2.63^, Trp75^2.64^, Pro92^3.32^, Phe96^3.36^, Phe161^ECL2^, Leu244^6.55^, Val2486^.59^, Ala250^6.61^, Pro259^7.32^, Leu262^7.35^, Leu263^7.36^, and Leu266^7.39^. Meanwhile, the terminal hydroxyl group forms a hydrogen bond with the polar residue Tyr240^6.51^ in TM6, known to play a critical role in GPBAR1 activation.[17,57–60] This interaction is particularly significant given that the presence of the 3-OH group in the α-configuration is a key determinant of the agonistic activity of bile acids.[60] The carboxylate tail, meanwhile, orients towards the extracellular vestibule, consistent with previously reported SAR data. Additional polar interactions are established with Tyr89^3.29^, Asn93^3.33^, Ser247^6.58^ and Tyr251^6.62^. Notably, this binding orientation closely mirrors that of INT-777 in the reference cryo-EM structure (RMSD = 0.08 nm, computed on the steroidal scaffold), further supporting the reliability of the docking results.

**Figure 2.**
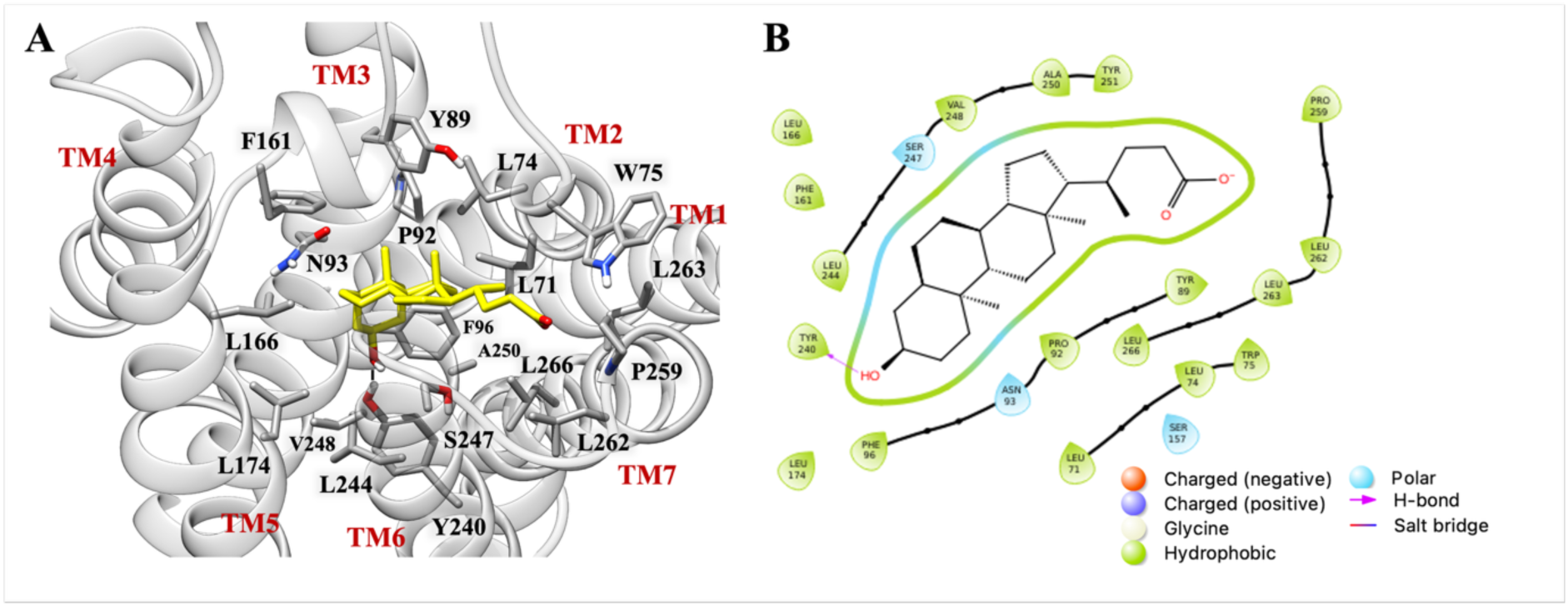
A) Docking pose of LCA in the GPBAR1-Gα*_s_*β_1_γ_2_ homology model. The ligand is represented as yellow sticks, while the interacting residues of the receptor are shown as gray sticks and labelled; oxygen and nitrogen atoms are colored red and blue, respectively. The receptor is represented as ribbons with its helices labelled. H-bonds are displayed as black dashed lines. B) Ligand Interaction Diagram (LID) of LCA in the GPBAR1-Gα*_s_*β_1_γ_2_ structural model.

To evaluate the stability and the energetics of the predicted LCA pose and its impact on the conformational dynamics of GPBAR1, the ligand-bound GPBAR1-Gα*_s_*β_1_γ_2_ complex was submitted to three independent 500 ns-long molecular dynamics (MD) simulations replica, in parallel with the ligand-free receptor-G protein assembly, thereby enhancing conformational sampling and improving statistical robustness. The inclusion of the G protein in both systems ensured that ligand-free and ligand-bound receptors could be compared from structurally and functionally equivalent starting points. Each system was embedded in an explicit 1-palmitoyl-2-oleoyl-sn-glycero-3-phosphocholine (POPC) and cholesterol (CHL) (molar ratio of 7:3) lipid bilayer, using physiological ionic conditions, to reproduce a native membrane environment.

Throughout the holo simulations, the LCA remained stably accommodated within the orthosteric pocket (Figure 3), with a slight deviation from the initial docking pose (average RMSD equal to 0.19 nm computed on the ligand’s heavy atoms) (Figure S1, panel S). This limited deviation indicates that the docking pose represents an energetically favorable binding mode that is preserved under dynamic conditions, supporting the robustness of the initial docking predictions.

**Figure 3.**
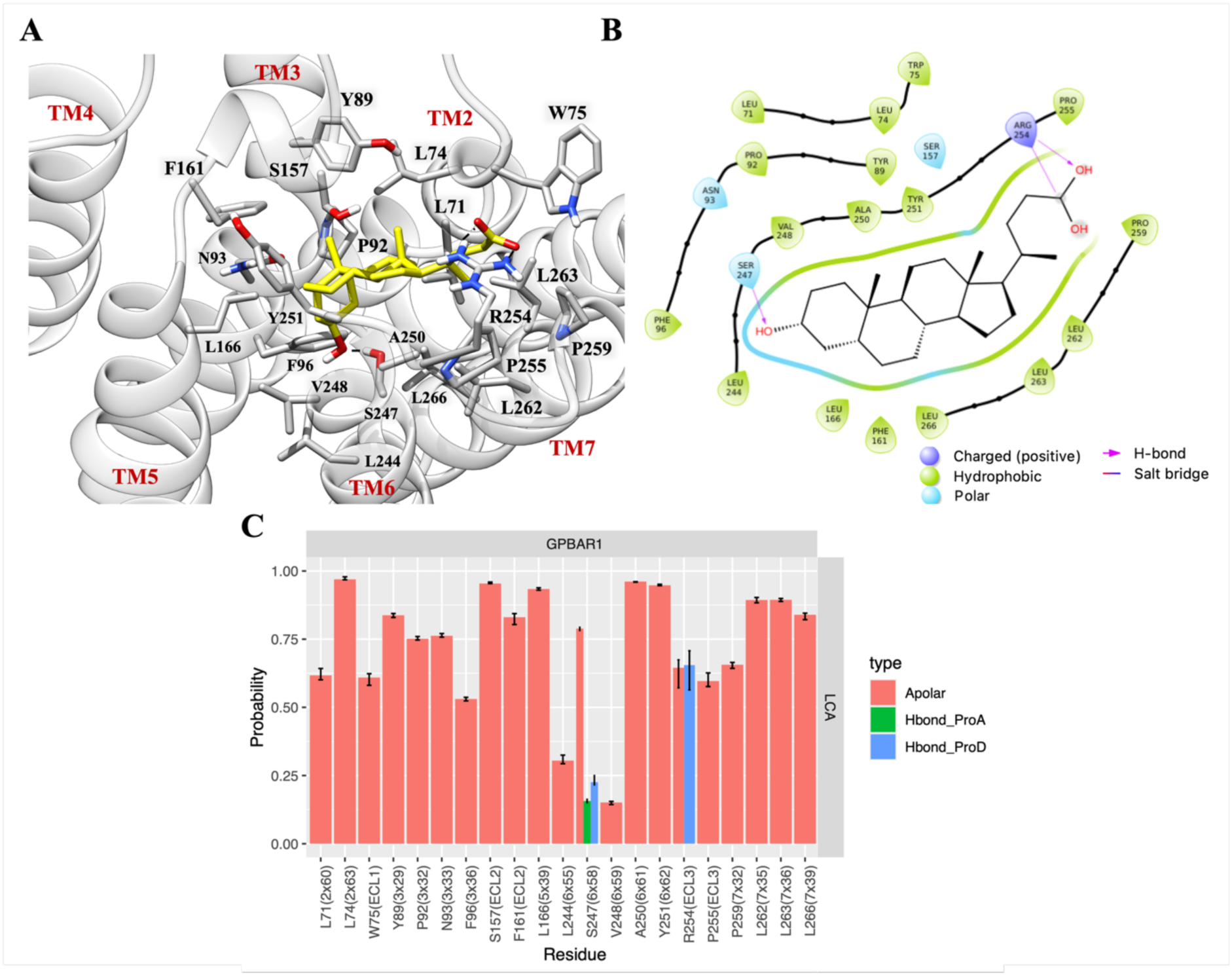
A) Centroid conformation of the most populated cluster obtained from the three MD replicas of the LCA in the GPBAR1-Gα*_s_*β_1_γ_2_ complex. The ligand is represented as yellow sticks, while interacting residues of the receptor are shown in gray and labeled; oxygen and nitrogen atoms are colored red and blue, respectively. The receptor is shown as ribbons with helices labeled. H-bonds are displayed as black dashed lines. B) Ligand Interaction Diagram (LID) of the MD-derived centroid of the most populated LCA binding mode in the GPBAR1-Gα*_s_*β_1_γ_2_ model. C) SIFt analyses showing probabilities of ligand-receptor interactions formed during three 500-ns MD simulations of the LCA-GPBAR1-Gα*_s_*β_1_γ_2_ system. Interaction types include apolar carbon-carbon contacts (Apolar, pink), hydrogen bond with the protein as acceptor (Hbond_ProA, green), and hydrogen bond with the protein as donor (Hbond_ProD, blue). Only interactions with an average probability greater than 10% are displayed.

The hydrophobic network involving the steroidal scaffold and residues in TM3, TM5, TM6 and the extracellular loops is largely preserved, maintaining the integrity of the ligand-binding cavity. Particularly, the ligand’s core interacts with Leu71^2.60^, Leu74^2.63^, Trp75^2.64^, Tyr89^3.29^, Pro92^3.32^, Asn93^3.33^, Phe96^3.36^, Ser157^ECL2^, Phe161^ECL2^, Leu166^5.39^, Leu244^6.55^, Val248^6.59^, Ala250^6.61^, Tyr251^6.62^, Pro255^ECL3^, Pro259^7.32^, Leu262^7.35^, Leu263^7.36^, and Leu266^7.39^.

Consistent with docking results, the MD-derived binding poses align well with site-directed mutagenesis and SAR data, which identify several of these residues, including Asn93^3.33^, Leu244^6.55^, Leu263^7.36^, and Leu266^7.39^, as key determinants of bile acid recognition and GPBAR1 activation.[59,61,62] Mutations at these positions have been shown to markedly reduce ligand affinity and agonist efficacy, supporting their direct involvement in orthosteric recognition and signaling transduction.[21]

Together, these residues participate in an extended hydrophobic interaction network that stabilizes LCA in a binding mode consistent with known functional determinants of GPBAR1, thereby reinforcing the mechanistic interpretation of the docking and MD results within the context of existing experimental evidence.[41,57,58,60,63–65]

In addition to transmembrane helices, MD simulations reveal a significant contribution of extracellular loop (ECL) residues to ligand stabilization. In particular, as confirmed by the structural interaction fingerprint (SIFt) analysis, (Figure 4, panel C) polar interactions between the LCA carboxylate group and Arg254^ECL3^, together with hydrogen bonding to Ser247^6.58^, suggest that extracellular regions contribute not only to ligand recognition but also to the dynamic stabilization of the bound agonist. Taken together, these data indicate that ECLs shape the ligand-binding vestibule and regulate access to the orthosteric site as observed in other class A GPCRs, [66] acting as a dynamic gate that stabilizes ligand residence and enhances receptor engagement and signaling efficacy.

**Figure 4.**
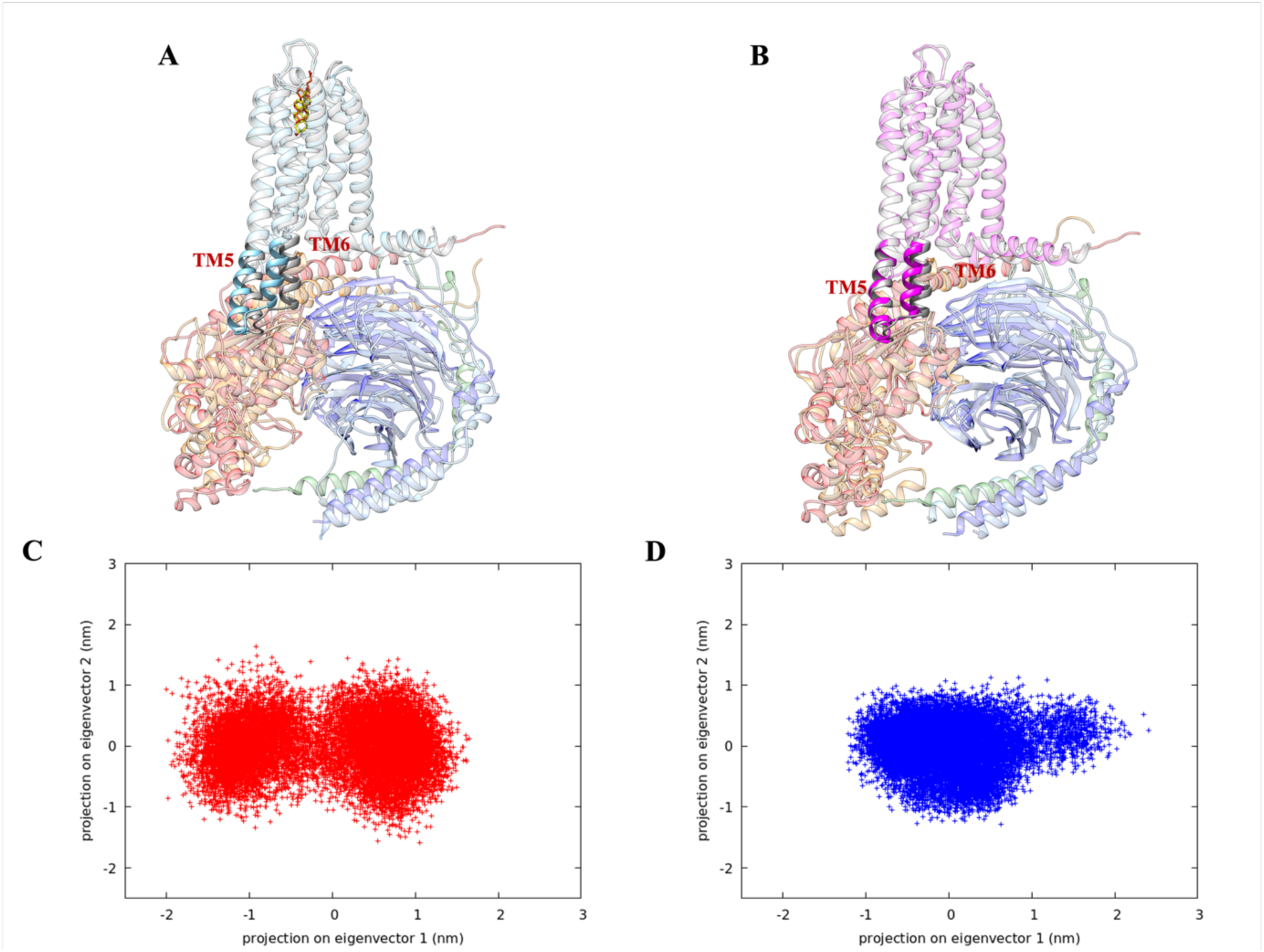
Superposition of the GPBAR1-G*_s_* model and MD-representative structures extracted from the most populated clusters for (A) the holo and (B) the ligand-free systems. The GPBAR1 model is shown in grey, while the MD-derived conformations are depicted in cyan for the holo system and in magenta for the ligand-free system. 2D projection of the first two principal components (PC1 and PC2), computed from PCA of the C_α_ atoms of TM5 and TM6, showing the conformational distributions of the (C) holo and (D) ligand-free systems.

### 3.3 LCA Binding Favors an Active-Like Conformation of GPBAR1

Comparative analysis of the ligand-free and holo trajectories indicates that LCA binding biases GPBAR1 conformational ensemble towards active-like states through coordinated rearrangements of TM5, TM6, and key activation microswitches. To characterize how ligand binding reshapes the receptor conformational landscape, we performed a Principal Component Analysis (PCA) on the C_α_ atoms of the receptor.

In the holo system, the conformational landscape spanned a broader region of Principal Component (PC) space (Figure 4, panel C and D), with the dominant collective motions involving an outward displacement of TM6 together with coordinated rearrangements of TM5. These motions are associated with the opening of the intracellular side of the receptor (Figure 4, panel A; Movie S1). The first principal component captures the largest collective motions involving TM5 and TM6, which are sampled with distinct amplitudes and directions in the holo *versus* ligand-free systems. By contrast, the ligand-free system explores a more restricted conformational space, populating a compact region of the PC space corresponding to conformations in which TM5 and TM6 remain more inward and closer to inactive-to-intermediate arrangements of the receptor (Figure 4, panel B).

Consistent with the PCA results, analysis of transmembrane helix rearrangements further supports a ligand-dependent reorganization of the intracellular region of GPBAR1. In the holo system, TM5 and TM6 undergo a more pronounced and persistent outward displacement that is consistently maintained across all replicas, as evidenced by RMSD/RMSF analyses (Figure S1, panels B, D, F, H, J; Figure S4). This observation is further supported by quantitative analysis of diagnostic Cα distances between residues located on TM3 and TM6, identified by Yang, *et al.*, [21] which revealed an increase in the separation of two helices in the holo system compared to the ligand-free receptor (Figure 5). Notably, the distribution of TM3–TM6 distances in holo GPBAR1 is shifted toward larger values compared to the ligand-free receptor, indicating a more pronounced intracellular opening. Although the magnitude of this separation is larger in GPBAR1 than in canonical class A GPCRs, active-state structures of the A_2_ adenosine receptor (A_2_AR) (PDB ID: 6GDG) and the β_2_-adrenergic receptor (β_2_-AR) (PDB ID: 3S6N) nonetheless display consistently wider TM3–TM6 distances relative to their corresponding inactive, G protein–free conformations (A_2_AR, PDB ID: 3VG9; β_2_-AR, PDB ID: 3YNA).

**Figure 5.**
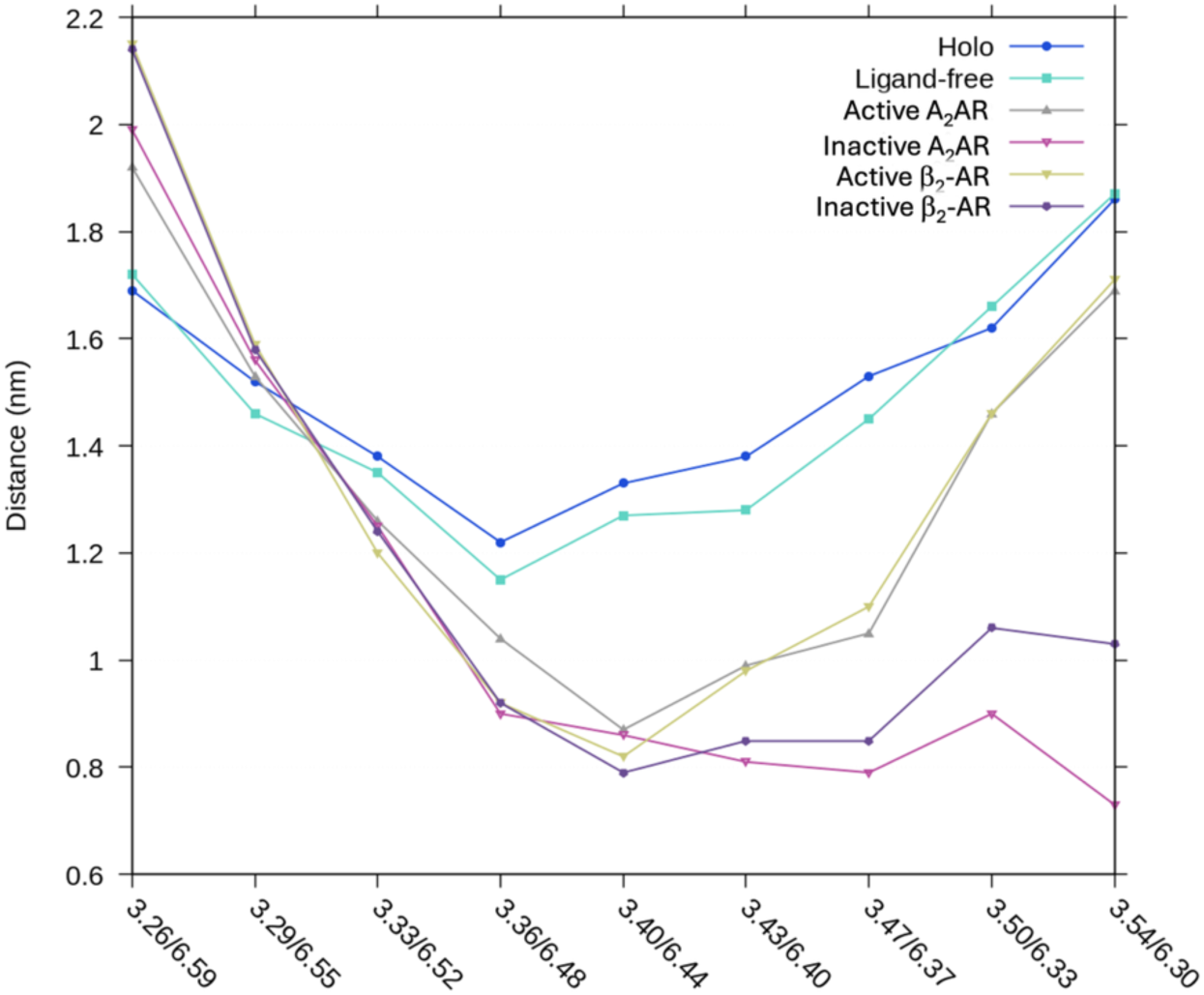
A) Plot of C_α_ distances between residues on TM3 and TM6 for the centroid of the most populated cluster of the holo and ligand-free GPBAR1 systems, alongside the active (PDB ID: 6GDG) and inactive (PDB ID: 3VG9) A_2_AR, and the active (PDB ID: 3S6N) and inactive (PDB: 3YNA) β_2_-AR. The x-axis shows the Ballesteros–Weinstein residue numbering.

By contrast, in the ligand-free simulation, GPBAR1 exhibited higher stability at the transmembrane bundle level, as reflected by the lower backbone RMSD values (Figure S1, panels A, C, E, G, I).

Additional support for ligand-induced stabilization of an active-like conformational ensemble was provided by conformational cluster analysis performed using the GROMOS algorithm.[48] In the ligand-free system, the trajectory is distributed across multiple conformational clusters, with the most populated cluster accounting for only 33% of the simulation frames, followed by several additional clusters with lower populations (Table S1). This broad distribution reflects increased conformational heterogeneity in the absence of ligand. In contrast, the holo system predominantly samples two closely related conformational clusters, representing 27% and 26% of the frames, respectively (Table S2). These clusters are structurally similar, with a backbone RMSD of 0.16 nm, indicating that LCA binding biases the conformational landscape of GPBAR1 toward a narrower basin centered around an active-like state while still allowing limited fluctuations between closely related conformations.

### 3.4 Ligand-Dependent Remodeling of Activation Microswitches and GPBAR1–Gα_s_ Interface

To investigate how the ensemble-level differences observed upon LCA binding translate into local activation features, we analyzed key receptor microswitches and intracellular motifs involved in G-protein coupling in both the ligand-bound and apo systems.

Trp^6.48^ is commonly regarded as the toggle switch in class A GPCRs, but previous reports have established that in GPBAR1 this functional role is primarily fulfilled by Tyr240^6.51^ rather than by Trp237^6.48^.[21] In agreement with this notion, analysis of the residue z-position along the membrane normal revealed that Tyr240^6.51^ occupies a lower position in the LCA-bound system compared to the ligand-free receptor (Figure S2, panel A). This downward displacement of Tyr240^6.51^ upon agonist binding highlights its central role as the primary sensor of ligand-induced activation in GPBAR1, while Trp237^6.48^ appears to participate in downstream conformational rearrangements within TM6, contributing to the stabilization of the active-like state rather than acting as the principal toggle switch (Figure S2, panel B).

GPBAR1 lacks the conserved NPxxY motif found in many class A GPCRs, while the canonical DRY motif is present as an ERY sequence.[21] To assess the functional relevance of this motif in the context of G protein coupling, we monitored throughout the simulations the hydrogen bond established between Glu109^3.49^ and 291 of Gα_s_ in the reference cryo-EM structure (Figure S3). This interaction is preferentially formed and stabilized in the presence of LCA, whereas in the ligand-free system it is either transient or absent, indicating that agonist binding promotes productive engagement of the ERY motif with the G protein. In addition, the inactivating ionic lock involving Arg110^3.50^, commonly observed in class A GCPRs, is intrinsically disrupted in GPBAR1 due to the substitution of the conserved counterpart Glu^6.30^ with a threonine. This residue may be capable of engaging in hydrogen bonding with Arg^3.50^ in the inactive state of the receptor, although this possibility would require dedicated investigation.

Consistent with the microswitch analysis and the TM5/TM6 rearrangements identified by PCA, ligand-dependent conformational differences propagated toward the intracellular side of the receptor. In the holo system, the coordinated outward motion of TM5 and TM6 promotes an increased opening of the G protein–binding cavity, thereby facilitating interactions with the C-terminal α5 helix of Gα*_s_*. The enhanced separation between TM5 and TM6 was dynamically correlated with increased flexibility and structural deviations of the α5 helix and the α-helical domain of Gα_s_ (Figure S1, panels K-P), indicating increased conformational heterogeneity at the receptor–G protein interface. Such behavior suggests that LCA binding may allosterically contribute to the early stages of G-protein engagement. In contrast, the Gβ and Gγ subunits remained conformationally stable in both ligand-free and holo simulations, exhibiting only minor positional fluctuations (Figure S1, panels Q-T).

Representative structures extracted from the most populated clusters further highlight pronounced differences between the ligand-free and holo states at the intracellular G protein-coupling interface (Figure 5 A and B). The ligand-free system, which represents a pre-formed yet inactive complex, exhibited higher structural stability, as reflected by the RMSD and RMSF profiles (Figures S1, S3-S5). In this state, the α5 helix remains stably inserted into the receptor cavity, supported by a hydrophobic cleft formed by residues from TM3, TM5, and TM6 (Figure 5, panel B; Figure S5). In contrast, in the holo system (Figure 5, panel A; Figure S4), the increased flexibility of TM5 and TM6 occasionally facilitates the unraveling and partial disengagement of the α5 helix from the receptor core. To quantitatively characterize this behavior, the radius of gyration (Rg) of the α5 helix was calculated using the GROMACS analysis tool. As shown in Figure S6, the holo simulations exhibit slightly higher Rg values compared to the ligand-free simulations, indicative of an increased conformational spread of the α5 helix in the presence of LCA. This behavior is consistent with rearrangements reported for early stages of G protein activation in agonist-bound GPCR complexes.[67,68]

### 3.5 Ligand Binding Enhances Specific Allosteric Communication Pathways to the G Protein Interface

To better discriminate the structural and dynamic differences between the holo and ligand-free states of the human bile acid receptor GPBAR1, we conducted a network-based analysis of allosteric communication pathways using the MDpath tool.[54] This approach infers residue-level communication routes by quantifying correlated atomic fluctuations along molecular dynamics trajectories and modeling them as probabilistic paths within a graph representation of the receptor (see Methods for details).

This analysis identified the most probable communication routes connecting the transmembrane (TM) helices and the intracellular region involved in G protein coupling. In both systems, the detected pathways are largely confined to the helical core of the receptor; however, clear differences emerged between the ligand-free and holo states. In the ligand-free system, the residues most involved in signal propagation are primarily located in TM5, whereas in the ligand-bound system they mainly belong to TM6, indicating a distinct redistribution of allosteric communication pathways between the two conformational ensembles (Figure S7). In the absence of ligand, dominant communication paths were shorter and more confined around the central bundle, particularly involving TM3-TM6 (Figure 6, panel A; Figure S7, panels A and B). These compact intrahelical networks are consistent with a more constrained intracellular region and with the partially relaxed active-like ensembles sampled in the ligand-free simulations, in which microswitch arrangements appear insufficient to support stable signal propagation toward the G protein coupling interface. [69,70]

**Figure 6.**
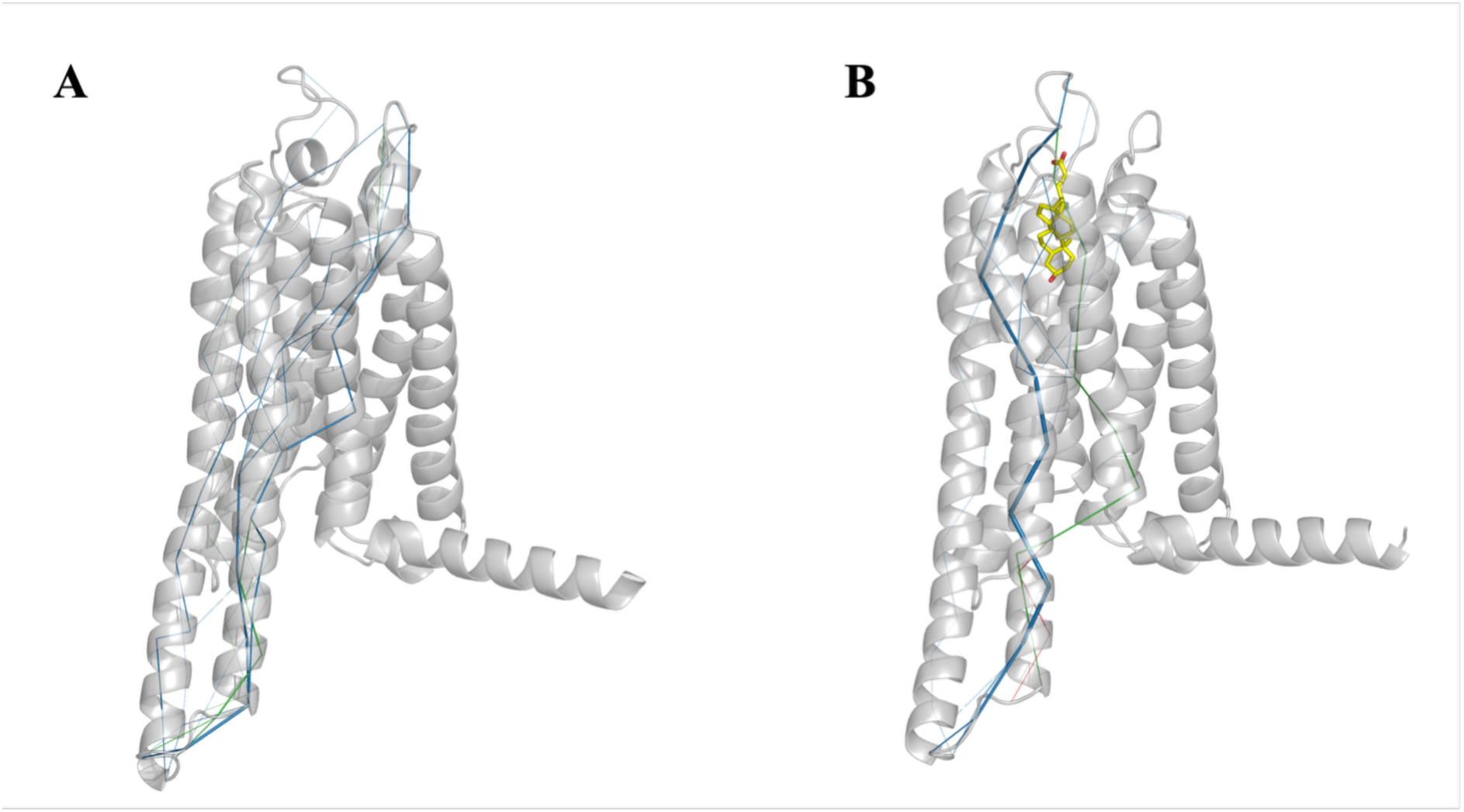
Comparison of allosteric communication pathways in GPBAR1 obtained from the MDpath tool. The ligand-free form (panel A) displays expanded communication paths, involving TM3-6, whereas the LCA-bound receptor (panel B) exhibits a more compact network centered on TM6, which is consistent with an active-like conformation. Communication paths are mapped onto backbone Cα, providing an intuitive representation of the significance of the residue-level allosteric network. Path colors correspond to the cluster assignments, enabling the different communication channels to be distinguished at a glance.

In contrast, LCA binding induces a redistribution of the communication routes towards TM6, reaching the intracellular region. This behavior correlates with the wider TM5-TM6 separation relative to TM3 discussed before and suggests enhanced dynamic coupling between the orthosteric pocket and the G protein coupling interface (Figure 6, panel B; Figure S7, panels A and C). In this context, ligand-induced stabilization of activation-associated microswitches and concerted TM5–TM6 rearrangements provides a structural framework that enables the emergence of extended allosteric communication across the receptor core.

Visual inspection of representative structures confirmed these findings: in the holo state, GPBAR1 (Figure 6, panel B; Figure S8) exhibits a pronounced outward movement of TM6 and a slight rotation compared to the ligand-free receptor (Figure 6, panel A; Figure S8), resulting in a more open intracellular cavity. Consistently, the dominant allosteric paths in the holo form aligned with this movement, running from the orthosteric site towards the cytoplasmic region along TM3-TM5-TM6.

Altogether, these results highlight how ligand binding reshapes the internal dynamic network of GPBAR1, facilitating the development of extended allosteric pathways that connect the extracellular and intracellular domains and preparing the receptor for G protein engagement.

## 4. Discussion

In this work, we present an integrative computational strategy aimed at exploring how LCA modulates the conformational landscape of GPBAR1 and its coupling with the G*_s_* protein. Although the available cryo-EM structures of the receptor provide a valuable framework,[21,71–73] it omits large flexible segments and does not capture the dynamic reorganizations that govern receptor signaling. By reconstructing the full receptor–G protein complex and comparing ligand-free and holo simulations, we investigated how LCA binding influences both local interactions within the orthosteric pocket and broader conformational features associated with receptor activation.

The ligand-binding mode obtained from docking (Figure 2) and subsequently refined using MD simulations (Figure 3) highlights how the endogenous bile acid leverages both hydrophobic packing and specific polar contacts within the transmembrane cavity. Importantly, these interactions are dynamic: throughout the MD simulations, the LCA hydroxyl group alternates between forming hydrogen bonds with Tyr240^6.51^ and Ser247^6.58^, while the carboxylate transiently establishes ionic contacts with Arg254^ECL3^. This behavior is captured in the centroid pose derived from clustering (Figure 3 and Figure S1, panel S). Such interaction patterns suggest that LCA accommodates intrinsic receptor flexibility instead of imposing a strictly rigid geometry, thereby modulating the receptor’s energy landscape rather than locking it into a unique conformation. Although no experimental structure of a GPBAR1–LCA complex is currently available, the present simulations provide an atomistic model that may help rationalize the high agonistic potency of this endogenous bile acid. In this context, the involvement of Ser247^6.58^ and Arg254^ECL3^ may contribute both to ligand stabilization and to the modulation of extracellular loop dynamics, which are increasingly recognized as integral components of GPCR activation networks.[66,74]

Beyond the binding pocket, LCA engagement is associated with a redistribution of intramolecular strain and with changes in the behavior of activation-related microswitches. The observed shift in the z-position of Tyr^6.51^ (Figure S2, panel A) and the outward displacement of TM6 are consistent with conformational features commonly associated with class A GPCR activation. These trends are in qualitative agreement with experimental observations reported by Yang *et al*.,[21] who described coordinated microswitch rearrangements and TM6 outward motion upon agonist binding. In the case of GPBAR1, these changes appear particularly sensitive to the positioning of the ligand core, supporting the idea that steroidal agonists act as conformational “wedges” that facilitate the outward displacement of TM6. This interpretation is further supported by the RMSD and RMSF analysis (Figure S1, S4, and S5), which show increased mobility of the intracellular tips of TM5 and TM6 in the holo state, indicative of a redistribution of flexibility from the extracellular pocket toward the intracellular interface.

The consequences of these reorganizations are clearly visible when comparing the conformational ensembles sampled by the holo and ligand-free systems. Clustering analysis indicates that the holo receptor explores a narrower subset of conformations (Figure 5, panel A), whereas the ligand-free system populates a more heterogeneous ensemble (Figure 5, panel B). PCA further reinforces this interpretation: the dominant global motion captures the outward displacement of TM6, which is prominently sampled in the holo simulations (Figure 5, panel C) but remains limited in the ligand-free state (Figure 5, panel D). Notably, while the holo receptor explores multiple active-like geometries, these conformations share a conserved TM6 displacement, suggesting that ligand binding biases the ensemble toward activation-competent states rather than stabilizing a single active structure. Conversely, the ligand-free receptor samples partially relaxed conformations characterized by reduced TM5–TM6 separation, consistent with a less efficiently coupled intracellular domain.

Additional insight emerges by examining the dynamics of the G protein, particularly the C-terminal α5 helix of Gα*_s_*. In the holo simulations (Figures S3–S5), the α5 helix exhibits enhanced fluctuations and occasional reductions in its depth of insertion into the intracellular cavity. These changes appear to correlate with the ligand-induced rearrangements of TM5 and TM6 and suggest a dynamic coupling between receptor activation and G protein conformational variability. Transient partial unraveling of α5 and rearrangements are observed within the nucleotide-binding site, particularly at the Ras–AHD interdomain region, which has been associated in previous studies with early stages of G protein activation.[67,68] While processes such as GDP destabilization and nucleotide exchange cannot be directly resolved on the timescale of the present simulations, the observed conformational trends are compatible with initial priming events that precede these steps and provide a structural context for interpreting how receptor activation may influence G protein readiness.

The network-based analysis of allosteric communication pathways further integrates these observations. In the absence of a ligand, communication pathways remain confined to the core of the TM bundle, particularly to the TM3–TM4 region, and do not effectively connect the orthosteric pocket to the intracellular interface. This suggests that ligand-free GPBAR1 lacks an efficient conduit for transmitting structural changes towards the G protein. Upon LCA binding, the dominant pathways redistribute towards TM5–TM6 and extend deep into the intracellular cavity. This reorganization supports the notion that ligand engagement reshapes the internal communication topology of GPBAR1, favoring pathways that connect extracellular ligand binding to intracellular signaling regions. The concentration of allosteric flow along TM6 in the holo state mirrors the structural expansion captured by PCA, highlighting the central role of this helix as a conduit for transmitting ligand-induced rearrangements. These features are consistent with experimentally described microswitch reconfigurations and suggest that the MD simulations capture not only ligand-induced local changes but also global structural propagation critical for G protein activation.[21,75] Altogether, the integration of structural modelling, atomistic simulations, and network analysis shows that GPBAR1 activation emerges from distributed conformational changes rather than a single structural transition. LCA binding stabilizes not one isolated active state, but a dynamically restricted ensemble optimized for G protein engagement, while retaining sufficient flexibility to respond to membrane composition, loop motions, and shifts in G protein dynamics. Although the present MD simulations capture dominant transitions associated with GPBAR1 activation, a complete mapping of the receptor’s metastable landscape will require enhanced-sampling methodologies and AI-assisted approaches, which are increasingly successful at resolving transient or experimentally inaccessible GPCR states that may prove crucial for ligand discovery and functional selectivity.[76,77]

In conclusion, this study refines our understanding of how steroidal ligands activate class A GPCRs by highlighting the importance of dynamic ensemble reshaping over static interaction patterns. These findings suggest that effective GPBAR1 modulators must be evaluated not only for their static interactions but also for their capacity to influence the receptor’s global dynamic architecture. The atomistic insights presented here provide a computational framework for future investigations into GPBAR1 pharmacology, and that may guide the rational design of next-generation ligands aimed at treating metabolic and inflammatory disorders.

## 5. Conclusion

In this study, we employed an integrative computational strategy combining structural modeling, molecular docking, and microsecond-scale molecular dynamics simulations to investigate the molecular and dynamic determinants of GPBAR1 activation by its endogenous agonist LCA. By reconstructing a full-length GPBAR1–G*_s_* assembly, we extended the information provided by available cryo-EM structures and explored receptor behavior beyond static conformations.

The simulations demonstrate that LCA binding stabilizes GPBAR1 toward a restricted ensemble of active-like conformations characterized by outward movements of TM5 and TM6 and by rearrangements of activation-related microswitches, including the GPBAR1-specific Tyr^6.51^ toggle switch. Rather than stabilizing a single active structure, ligand engagement reshapes the receptor’s dynamic landscape, favoring conformations compatible with intracellular cavity opening and G protein interaction.

At the orthosteric site, the dynamic binding mode of LCA highlights the contribution of transient polar contacts and a conserved hydrophobic network involving TM3, TM5, and TM6, along with extracellular loops. These features, not readily accessible from static experimental structures, provide additional insight into the molecular basis of ligand recognition and potency in GPBAR1 and may inform future structure–activity relationship studies.

Finally, analysis of allosteric communication pathways suggests that ligand binding reorganizes internal signaling routes, enhancing connectivity between the orthosteric pocket and the intracellular coupling interface. Together, these findings support a model in which GPBAR1 activation emerges from distributed, ligand-dependent conformational changes rather than from a single structural transition.

The integrative computational approach presented here establishes a foundation for future investigations into GPBAR1 pharmacology and provides key information for the rational development of next-generation modulators targeting metabolic and inflammatory diseases in which this receptor plays a central role.

## CRediT authorship contribution statement

**Bianca Fiorillo**: Conceptualization, Data curation, Formal analysis, Investigation, Methodology, Project administration. Validation, Visualization, Writing - original draft.

**Federica Moraca:** Writing - review & editing, Validation, Visualization.

**Valentina Sepe:** Writing - review & editing, Validation, Visualization.

**Francesco Saverio Di Leva:** Writing - review & editing, Validation, Visualization.

**Stefano Fiorucci:** Writing - review & editing, Validation, Visualization.

**Vittorio Limongelli:** Writing - review & editing, Validation, Visualization.

**Angela Zampella:** Writing - review & editing, Validation, Visualization.

**Bruno Catalanotti**: Conceptualization, Funding acquisition, Project administration. Resources, Software, Supervision, Writing - review & editing, Validation, Visualization.

## Appendix A. Supplementary data

Supplementary data to this article can be found online.

## Acknowledgements

This work was partially supported by a grant from the Italian MUR/PRIN 2022 PNRR (No. P20227JB3W). Vittorio Limongelli acknowledges funding from the European Research Council (ERC) under the European Union’s Horizon 2020 research and innovation programme (“CoMMBi” ERC grant agreement No. 101001784), the Swiss National Science Foundation (SNSF) (grant number IC00I0-231546), and the Swiss National Supercomputing Centre (CSCS) under project ID lp40 and u8.

## Abbreviations

GPBAR1: G protein-coupled bile acid receptor 1
TGR5: Takeda G protein-coupled receptor 5
AHD: α-helical domain
LCA: Lithocholic acid
SAR: Structure–activity relationship
TM: Transmembrane helix
ECL: Extracellular loop
MD: Molecular Dynamics
PCA: Principal Component Analysis
RMSD: Root mean square deviation
RMSF: Root mean square fluctuation
Rg: Radius of gyration
NMI: Normalized mutual information
SIFt: Structural interaction fingerprint
GUI: Graphical user interface
PRCG: Polak–Ribiere conjugate gradient
OPLS: Optimized potentials for liquid simulations
CGenFF: CHARMM General Force Field
DOPE: Discrete optimized protein energy
cryo-EM: Cryo-electron microscopy
PDB: Protein Data Bank
PBC: Periodic boundary conditions
POPC: 1-palmitoyl-2-oleoyl-sn-glycero-3-phosphocholine
CHL: Cholesterol
GlcNAc: N-acetylglucosamine
cAMP: Cyclic adenosine monophosphate
PKA: Protein kinase A
GLP-1: Glucagon-like peptide-1
IBD: Inflammatory bowel disease.

## Data Availability

The data that support the findings of this study are openly available in Zenodo at www.zenodo.org, reference DOI: 10.5281/zenodo.17883284.

