## Supplemental for "From Structure to Dynamics: Activation Mechanism of the G Protein-Coupled Bile Acid Receptor 1-G*s* Complex"

### Table of contents

#### Supplemental Figures and Figure Legends

**Figure S1:** Average RMSD plots of the three 500ns-long MD replicas. **S2**
**Figure S2:** Violin plot showing the vertical displacement (z-position) of the toggle switches Tyr$^{6.51}$ and Trp$^{6.48}$. **S3**
**Figure S3:** Time evolution of the hydrogen-bond distance between Glu109$^{3.49}$ of the ERY motif in GPBAR1 and Tyr291 of the G$\alpha_s$ $\alpha$5 helix over three independent MD simulation replicas. **S3**
**Figure S4:** Root Mean Square Fluctuation (RMSF) analysis of the LCA–GPBAR1–G$_s$ system computed over three independent 500ns MD replicas. **S4**
**Figure S5:** Root Mean Square Fluctuation (RMSF) analysis of the ligand-free GPBAR1–G$_s$ system computed over three independent 500ns MD replicas. **S4**
**Figure S6:** Radius of gyration (Rg) computed on the G$\alpha_s$ $\alpha$5-helix backbone over three independent replicas (500 ns each). **S5**
**Figure S7:** Allosteric communication pathways in GPBAR1 obtained from MDpath analysis. **S6**
**Figure S8:** Radius of gyration (Rg) computed on the GPBAR1 receptor TM6 backbone over three independent replicas (500 ns each). **S7**

#### Supplemental Tables

**Table S1:** Cluster analysis of 1500 ns MDs of ligand-free GPBAR1–G protein system. **S8**
**Table S2:** Cluster analysis of 1500 ns MDs of LCA-bound GPBAR1–G protein system. **S8**

#### Supplemental Movie

**Movie S1:** Principal component analysis (PCA) of GPBAR1–G$\alpha_s\beta_1\gamma_2$ systems.

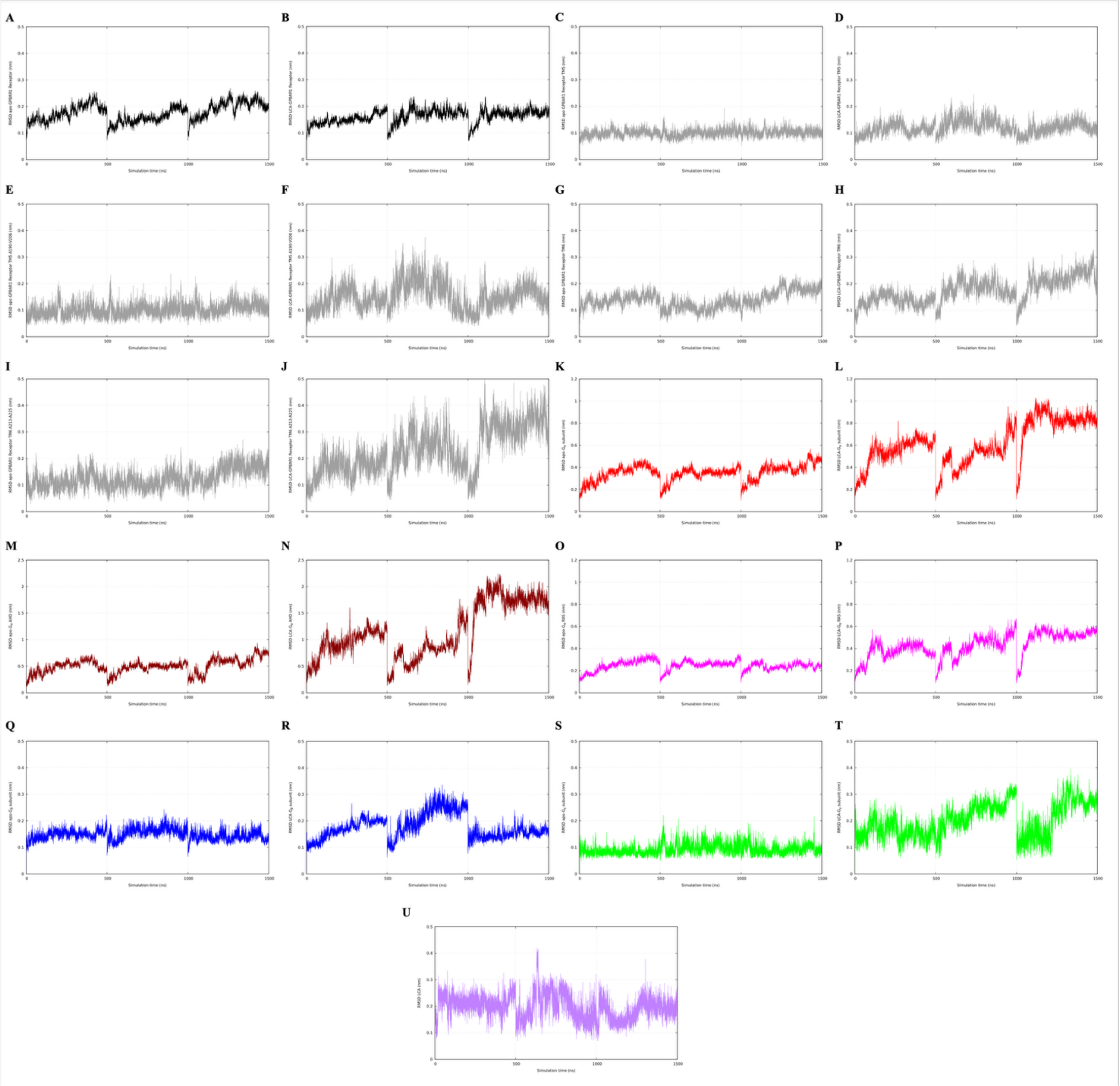


**Figure S1.** Average RMSD plots of the three 500 ns-long MD replicas calculated for: A–B) the receptor backbone in the ligand-free and holo states, respectively; C–D) the TM5 backbone in the ligand-free and holo states, respectively; E–F) the intracellular portion of TM5 in the ligand-free and holo states, respectively; G–H) the TM6 backbone in the ligand-free and holo states, respectively; I–J) the intracellular portion of TM6 in the ligand-free and holo states, respectively; K–L) the Gα subunit backbone in the ligand-free and holo states, respectively; M–N) the AHD backbone in the ligand-free and holo states, respectively; O–P) the RAS domain backbone in the ligand-free and holo states, respectively; Q-R) the Gβ subunit backbone in the ligand-free and holo states, respectively; S-T) the Gγ subunit backbone in the ligand-free and holo states, respectively; U) the ligand heavy atoms.

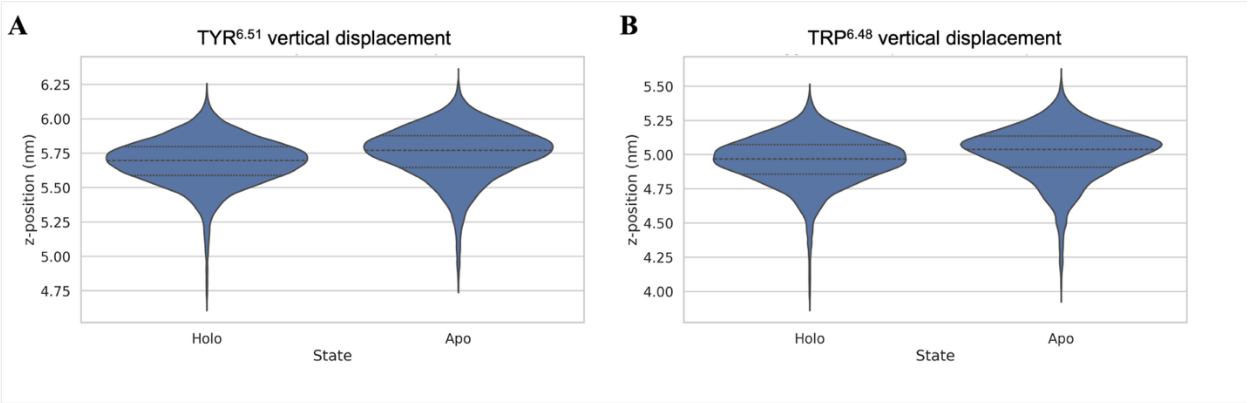


**Figure S2.** Violin plot showing the vertical displacement (z-position) of A) the toggle switches Tyr$^{6.51}$ and B) Trp$^{6.48}$, calculated as the center-of-mass position of each residue in both ligand-free and ligand-bound (holo) systems across the MD simulation. Positional values are reported in nanometers (nm).

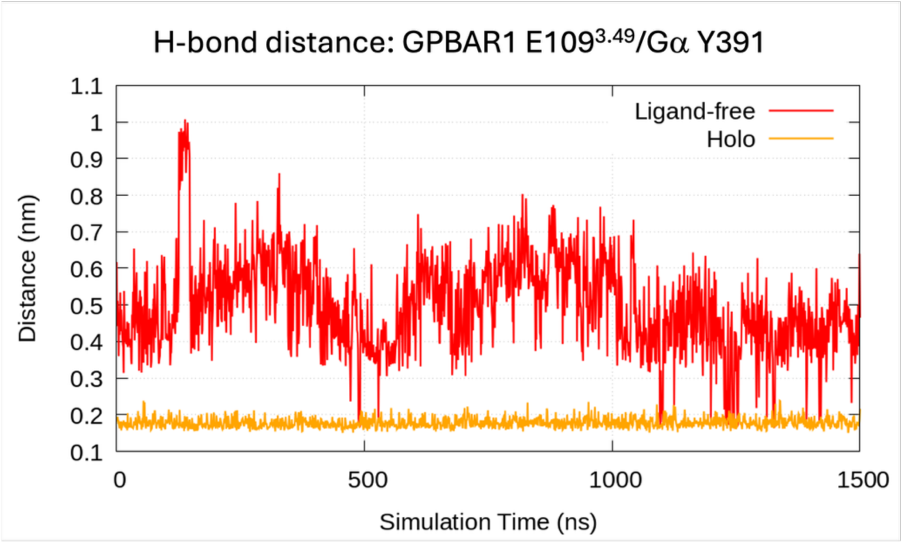


**Figure S3.** Time evolution of the hydrogen-bond distance between Glu109$^{3.49}$ of the ERY motif in GPBAR1 and Tyr291 of the G$\alpha_s$ α5 helix over three independent MD simulation replicas. Distances are shown for both the ligand-free and LCA-bound (holo) systems, highlighting that the interaction is maintained predominantly in the holo complex, indicative of ligand-stabilized coupling between the receptor and G protein.

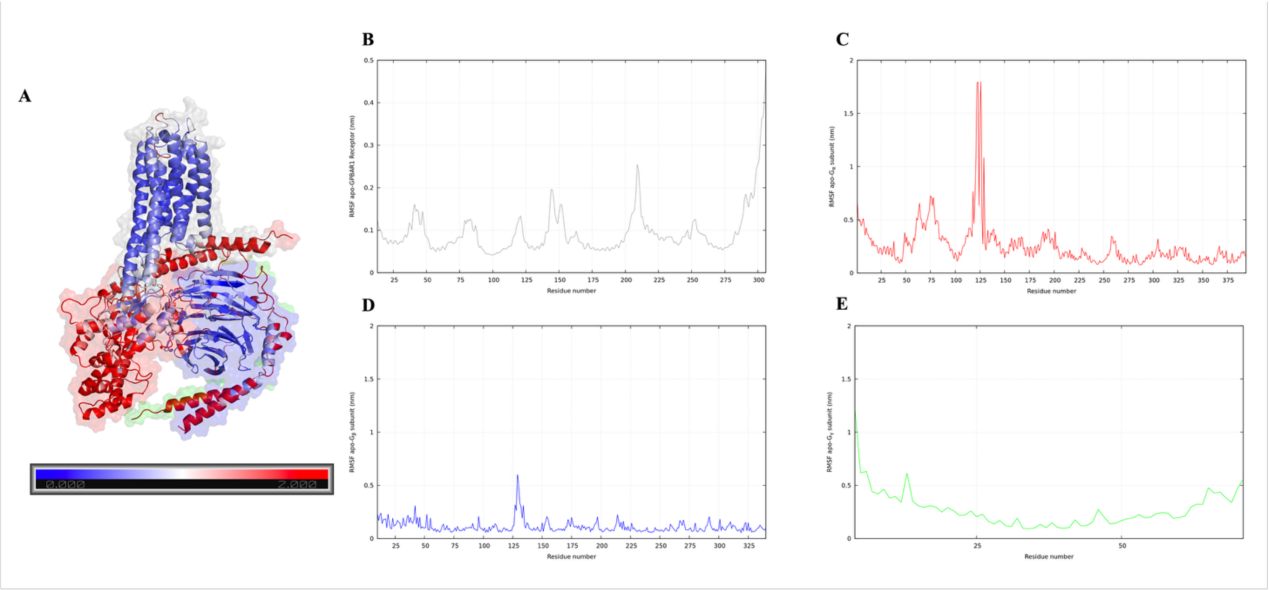


**Figure S4.** Root Mean Square Fluctuation (RMSF) analysis of the LCA–GPBAR1–G$_s$ system computed over three independent 500ns MD replicas. A) Structural representation of the LCA–GPBAR1–G$_s$ complex colored according to the averaged RMSF-derived B-factors (scale 0–2). The intensity of the colors reflects the local structural flexibility: blue tones indicate low mobility, white intermediate mobility, and red tones high mobility of the backbone atoms; LCA is depicted as yellow spheres, with oxygen atoms in red and nitrogen in blue. Panels B–E show the average RMSF profiles for: B) the GPBAR1 backbone; C) the Gα subunit backbone; D) the Gβ subunit backbone; and E) the Gγ subunit backbone. Higher RMSF values denote regions of increased structural mobility across the complex.

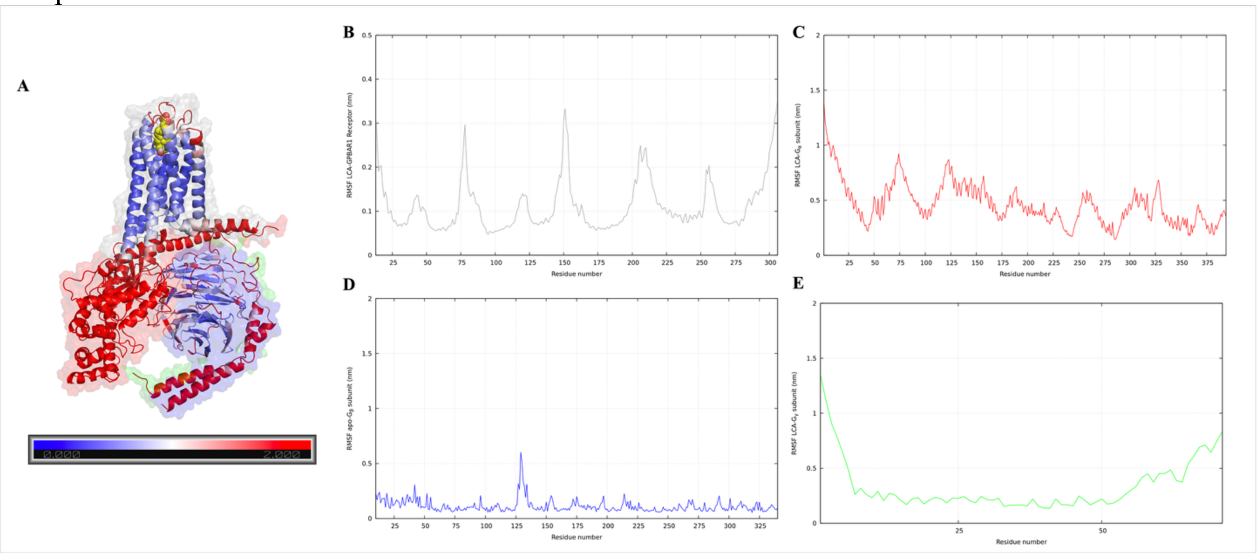


**Figure S5.** Root Mean Square Fluctuation (RMSF) analysis of the ligand-free GPBAR1–G$_s$ system computed over three independent 500ns MD replicas. A) Structural representation of the ligand-free GPBAR1–G$_s$ complex colored according to the averaged RMSF-derived B-factors (scale 0–2). The intensity of the colors reflects the local structural flexibility: blue tones indicate low mobility, white intermediate mobility, and red tones high mobility of the backbone atoms. Panels B–E show the average RMSF profiles for: B) the GPBAR1 backbone; C) the Gα subunit backbone; D) the Gβ subunit backbone; and E) the Gγ subunit backbone. Higher RMSF values denote regions of increased structural mobility across the complex.

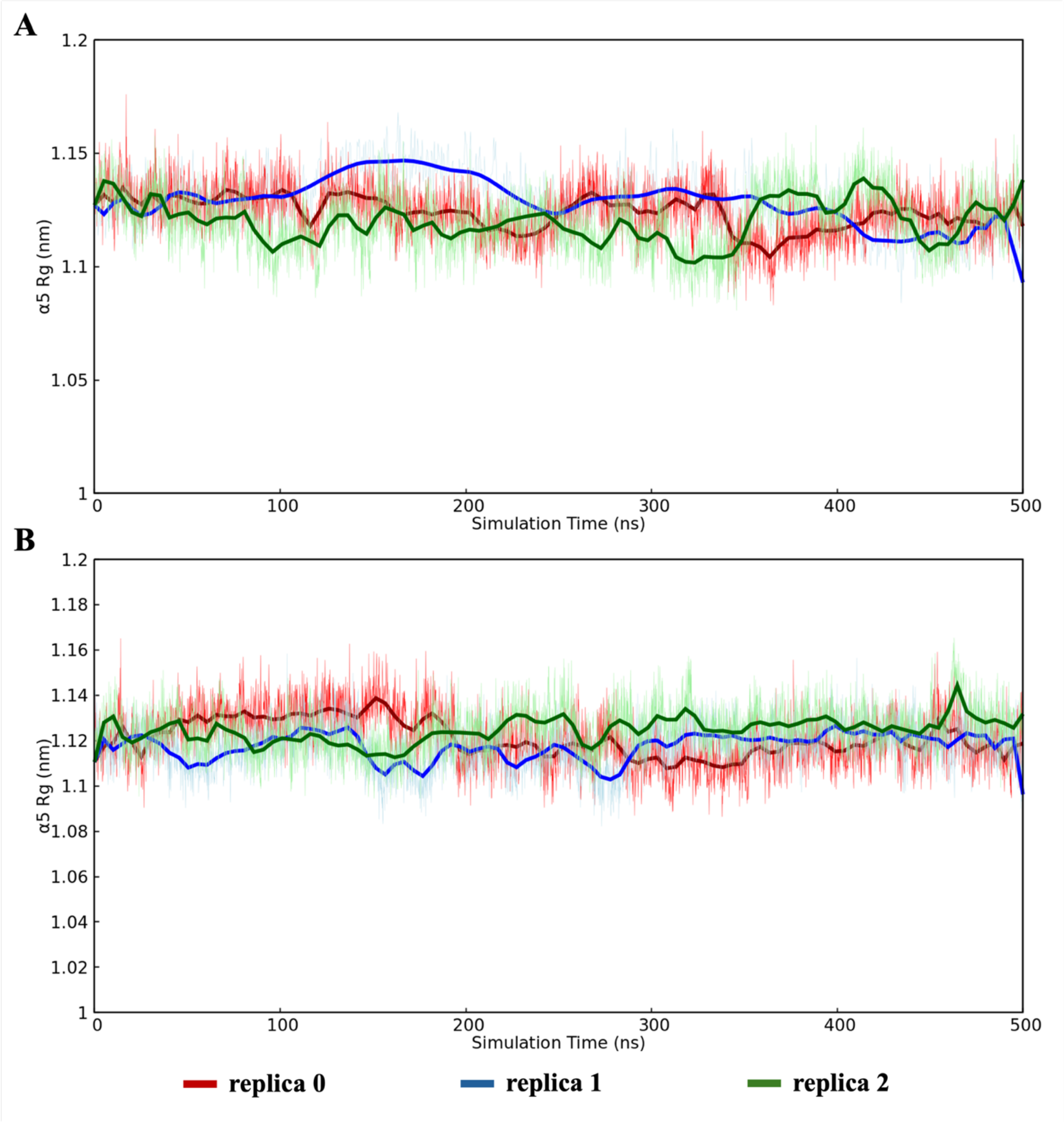


**Figure S6.** Radius of gyration (Rg) computed on the G$\alpha_s$ α5-helix backbone over three independent replicas (500 ns each) in A) LCA– and B) ligand-free-GPBAR1–$G_s$ systems. The red, blue and green traces correspond to the replica 0, 1, and 2, respectively. Overall, both systems maintain a stable global compactness of the α5 helix throughout the simulations, with the holo complex displaying slightly enhanced fluctuations consistent with ligand-induced modulation of G-protein engagement and with transient partial unraveling and disengagement events of α5.

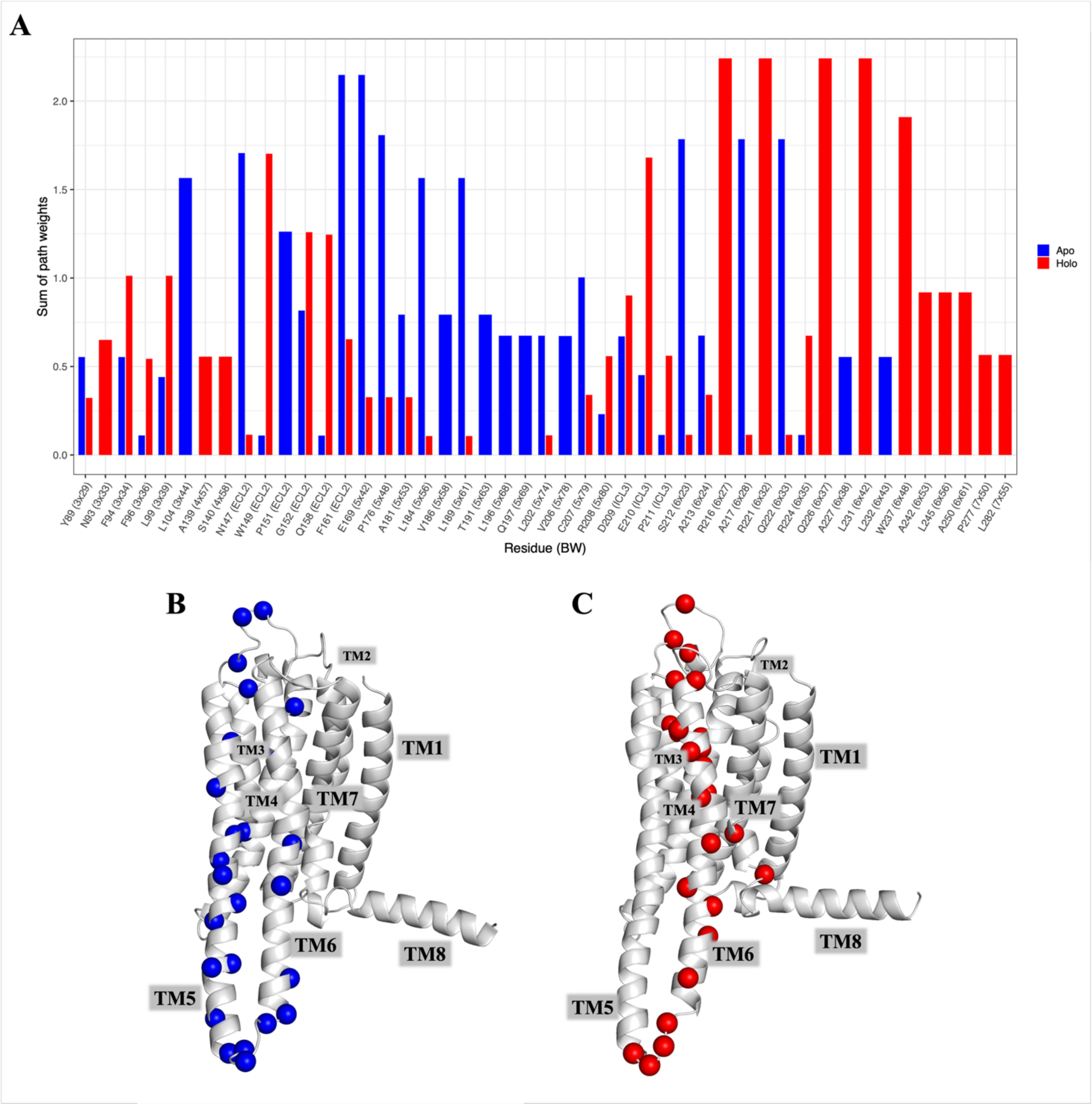


**Figure S7.** Allosteric communication pathways in GPBAR1 obtained from MDpath analysis. A) Plot of residues involved in allosteric communication; comparison of the ligand-free (panel B) and ligand-bound (panel C) states. Pathways are represented by Cα atoms represented as blue and red spheres for the ligand-free and ligand-bound systems, respectively.

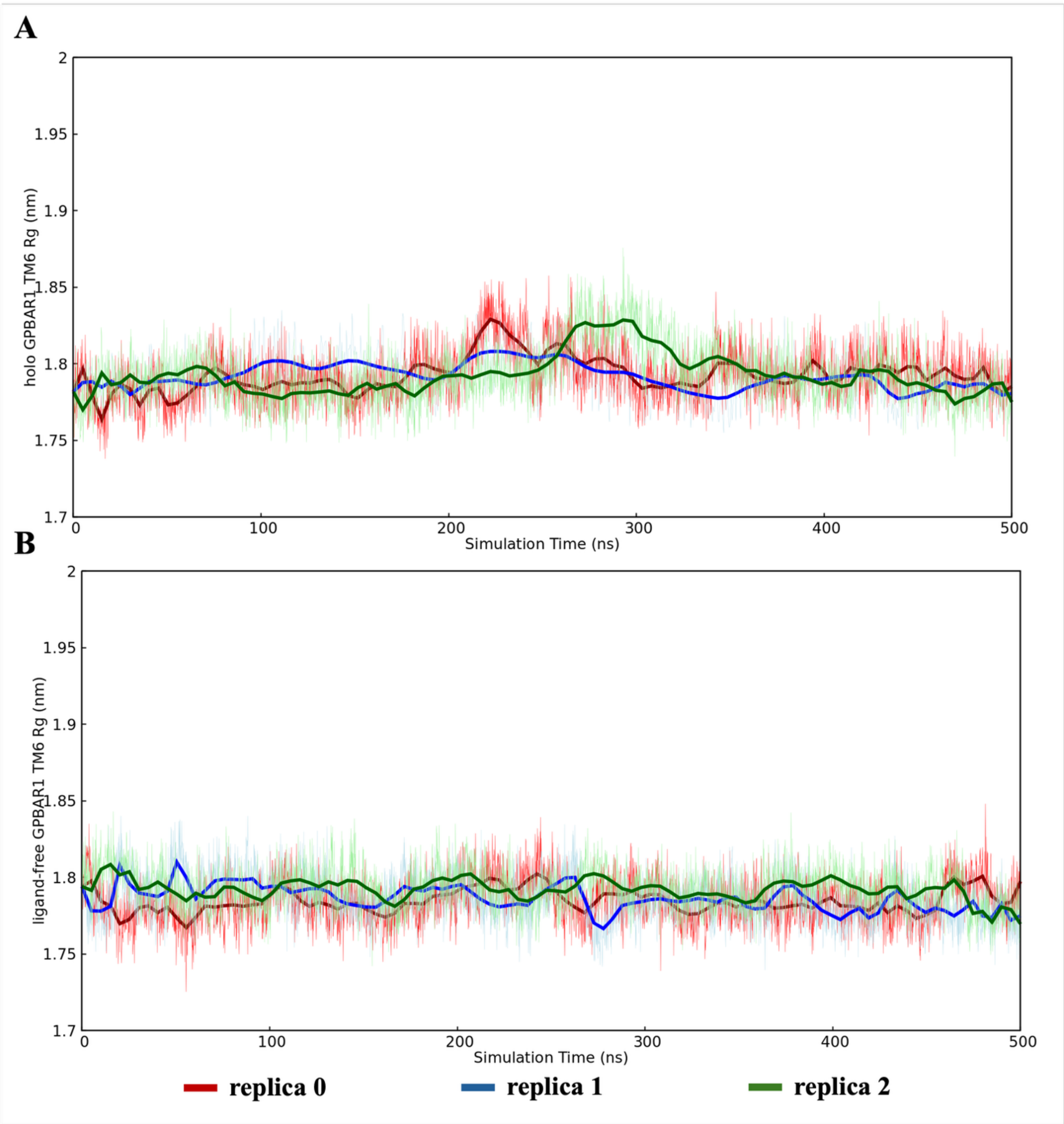


**Figure S8.** Radius of gyration (Rg) computed on the GPBAR1 receptor TM6 backbone over three independent replicas (500 ns each) in A) LCA– and B) ligand-free-GPBAR1–G$_s$ systems. The red, blue and green traces correspond to the replica 0, 1, and 2, respectively. Overall, both systems maintain a stable global compactness of the TM6 helix throughout the simulations, with the holo complex displaying slightly enhanced fluctuations consistent with ligand-induced modulation of the receptor activation.

**Table S1.** Cluster analysis of 1500 ns MDs of ligand-free GPBAR1–G protein system.

| Cluster number | % of population | Average rmsd (nm) |
|---|---|---|
| 1 | 33% | 0.299 |
| 2 | 24% | 0.287 |
| 3 | 13% | 0.293 |
| 4 | 11% | 0.304 |

**Table S2.** Cluster analysis of 1500 ns MDs of LCA-bound GPBAR1–G protein system. Notably, the first two clusters calculated for the holo system were structurally similar, with a backbone RMSD of 0.16 nm.

| Cluster number | % of population | Average rmsd (nm) |
|---|---|---|
| 1 | 27% | 0.300 |
| 2 | 26% | 0.287 |
| 3 | 10% | 0.314 |

**Movies S1.** Principal component analysis (PCA) of GPBAR1–G$\alpha_s\beta_1\gamma_2$ systems. The GPBAR1 receptor is shown in cyan cartoon, with TM5 and TM6 of the ligand-free system colored in blue, and the corresponding TM helices of the LCA-bound system in red. The movie illustrates the main conformational transitions associated or not with LCA binding.
